# Effect of crocin and naringenin supplementation in cryopreservation medium on post-thawed rooster sperm quality and expression of apoptosis associated genes

**DOI:** 10.1101/846758

**Authors:** Mahdieh Mehdipour, Hossein Daghigh Kia, Abouzar Najafi

**Author notes:** Corresponding author: Department of Animal Science, College of Agriculture, University of Tabriz, Tabriz, Iran. E-mail addresses, (H.D. Kia).

## Abstract

The aim of our research was to examine the effects of crocin (0.5 (C0.5), 1 (C1) and 1.5 (C1.5) mM) and naringenin (50 (N50), 100 (N100) and 150 (N150) µM) in cryopreservation extender for freezing rooster semen. Sperm motility, viability, abnormalities, membrane integrity, mitochondrial activity, apoptosis status, lipid peroxidation (LP), GPX, SOD, TAC, the mRNA expression of pro-apoptotic (CASPASE 3) and anti-apoptotic (Bcl-2) genes, fertility and hatchability rate were investigated following freeze-thawing. C1 and N100 resulted in the higher (P < 0.05) total motility and progressive motility in comparison to the control group. C1 and N100 improved viability, membrane integrity and reduced lipid peroxidation. We found much higher values for mitochondria activity with C1 and N100 respect to the control group. The C1 and N100 showed lower percentages of early apoptosis when compared with control group. Also, C1 and N100 had higher TAC when compared with control group. The mRNA expression of BCL-2 in the C1 and N100 group were significantly higher than that of other treatments. The expression of CASPASES 3 was significantly reduced in C1 and N100 group (P < 0.05) when compared to control group. Significantly higher percentage of fertility and hatching rate were observed in C1 and N100 compared to the control group. In conclusion, crocin at 1 mM and naringenin at 100 µM seem to improve the post-thawing rooster semen quality, fertility and could protect the sperm against excessive ROS generation by reducing the pro-apoptotic (CASPASE 3) and increasing anti-apoptotic (Bcl-2) genes.

## 1. Introduction

Despite its utilization over 70 years ago [1], cryopreservation of bird sperm causes low fertility, which limits its applying in genetic stock preservation [2]. Cryopreservation causes harmful effects on sperm that decrease sperm viability and motility [3–5]. Avian sperm are particularly susceptible to oxidative stress [6], though reactive oxygen species (ROS), in physiological quantities, are necessary for important sperm events leading to successful fertilization [7]. In sperm, oxidative stress disturb motility and mitochondrial activity [8]; induces lipid peroxidation of the membrane [9]; and the oxidation and DNA fragmentation [10].

Adding antioxidant compounds to the freezing medium is known as one of the ways to defeat the deleterious effects of ROS on sperm fertility after thawing, because it blocks or inhibits oxidative stress. Antioxidants provide a positive effect on semen, leading to an improvement in some sperm parameters containing motility and membrane integrity [11–13].

Naringenin is known as a natural flavonoid that has been studied for some of the most prominent properties containing antioxidant, antiproliferative, anti-inflammatory, and antimutagenic ones [14]. It was observed in previous experimental studies that naringain protects the cells from lead and arsenic-induced oxidative damage [15, 16].

The other studied antioxidant was crocin, a glycosyl ester of crocetin (one of the carotenoids extracted from saffron) [17]. In an experiment which was performed under in vitro conditions, crocin had an effect on improving deer sperm motility [18]. This antioxidant can influence sperm physiology through its protective effect on sperm cryopreservation media.

To the best of our knowledge, no similar study has been performed to evaluate the potential effect of naringenin and crocin in cryopreservation of rooster sperm. The objective of this investigation was to determine the effect of various levels of naringenin and crocin in the extender on post-thawed rooster sperm quality and expression of apoptosis associated genes. Quality and fertility analyses of the post-thaw sperm integrated with naringenin and crocin were also performed after the freezing and thawing process.

## 2. Materials and methods

### 2.1. Chemicals and ethics

All chemicals used for performing this experiment were purchased from Sigma (St. Louis, MO, USA) and Merck (Darmstadt, Germany) chemical companies. Approval for the present experiment was given by The Research Ethics Committees of the University of Tabriz.

### 2.2. Rooster and semen collection

This study was performed on ten adult Ross 308 broiler breeder roosters (30 week old) which were kept individually in cages (diet compositions were included: 12% crude protein and 2,750 kcal maintenance energy/kg). Semen was collected twice a week from individual birds in a graduated plastic tube [19]. Semen samples from each rooster were analyzed individually. The samples that had the standard criteria motility of >80% concentration of >3 × 10^9^ sperm/mL and volume of >0.2 mL were used in the present study. Next, to remove individual differences, semen samples were pooled and then assigned into 7 equal aliquots.

### 2.3. Extender preparation and cryopreservation

Seven experimental groups were applied in this study for semen dilution (Table 1): Beltsville extender without antioxidant (control), C0.5 (Beltsville extender with 0.5 mM crocin), C1 (Beltsville extender with 1 mM crocin), C1.5 (Beltsville extender with 1.5 mM crocin), N50 (Beltsville extender with 50 µM naringenin), N100 (Beltsville extender with 100 µM naringenin), N150 (Beltsville extender with 150 µM naringenin). Glycerol was added to the extender at 3.8% (v/v). Next, diluted semen samples were aspirated into 0.25 ml French straws (IMV, L’Aigle, France) to attain the concentration of 100 × 10^6^ sperm/mL. Consequently, via polyvinyl alcohol powder were sealed and equilibrated at 4 °C for 3 h. Then, after equilibration time (3 h), the straws were cryopreserved in liquid nitrogen (LN) vapor (4 cm above the LN for 7 min in a cryobox). Then, the straws were plunged into LN for storage until thawed (37 °C for 30 s) and used for assessment of sperm parameters.

**Table 1.** Composition of the Beltsville extender.

| Ingredients |  |
| --- | --- |
| Potassium citrate tribasic monohydrate (g) | 0.64 |
| Sodium-L-glutamate (g) | 8.67 |
| Magnesium chloride anhydrous (g) | 0.34 |
| D-(–)-Fructose (g) | 5 |
| Potassium phosphate dibasic trihydrate (g) | 7.59 |
| Potassium phosphate monobasic (g) | 0.7 |
| N-[Tris (hydroxymethyl) methyl]-2 (g) | 2.7 |
| Sodium acetate trihydrate (g) | 3.1 |
| Purified water (mL) | 100 |
| pH | 7.1 |
| Osmolality (mOsm/kg) | 310 |

### 2.4. Motility characteristics

Sperm motility and velocity parameters were determined using a computer-assisted sperm analyzer (CASA). To perform this, semen were diluted (1:10) by PBS buffer. Next, 10 μl of sperm sample was dropped onto a pre-warmed chamber slide (37 °C, Leja 4; Leja Products, Luzernestraat B.V., Holland). At least five fields containing a minimum of 200 sperm, were assessed by CASA. Sperm total motility (TM, %), progressive motility (PM, %), average path velocity (VAP, µm/s), straight linear velocity (VSL, µm/s), curvilinear velocity (VCL, µm/s), and amplitude of lateral head displacement (ALH, µm) were evaluated [3].

### 2.5. Viability

Sperm viability was evaluated by the eosin-nigrosine method described by Amini, Kohram (20). A 5 µl of sperm and 10 µl eosin-nigrosine stains was spread on a slide. To detect sperm viability, 200 sperm were assessed under a bright-field microscope at 400 ×.

### 2.6. Membrane integrity

Evaluating sperm membrane functionality was performed by Hypoosmotic swelling test (HOST) [21]. The assay was performed by adding 10 μL of diluted semen into eppendorf tubes containing 100 mL hypoosmotic solution (1.9 mM sodium citrate and 5 mM fructose, 100 mOsm/kg). After incubation at 37 °C for 30 min, total of 10 μL of the sample was poured on a microscope slide, and 200 sperm instantly was calculated under phase-contrast microscope at ×400 to detect sperm membrane integrity.

### 2.7. Morphology

For the assessment of morphology after thawing, 10 μL of sperm were pipetted into tubes including 1 ml of Hancock solution [22] (150 ml sodium saline solution, 150 ml PBS buffer solution and 62.5 ml formalin (37%)). To detect sperm total abnormality, about 200 sperm were counted by phase-contrast microscope at×1000.

### 2.8. Malondialdehyde (MDA) levels

MDA levels were assessed by thiobarbituric acid reaction [23]. In brief, 1 mL of sperm was mixed with 1 ml of cold trichloroacetic acid (20%) to precipitate protein. subsequently, the samples were centrifuged (963×g for 15 min), and 1 ml of the supernatant was incubated with tubes containing 1 ml of thiobarbituric acid (0.67%) in a boiling water bath at 100 °C for 10 min. After cooling, the absorbance was assessed by a Instruments Ltd, UK) at 532 nm.

### 2.9. TAC, GPx and SOD assessment

The antioxidant system was examined by assessment of GPx, TAC, and SOD levels [24]. This variable was assessed spectrophotometrically by Randox™ kits (RANDOX Laboratories Ltd.) and an Olympus AU 400 automatic biochemistry analyzer (Olympus, Tokyo, Japan).

### 2.10. Flow cytometry

Mitochondria activity and apoptosis status were analyzed by FACSCalibur flow cytometer (Becton Dickinson System, San Jose, CA, USA). The excitation wavelength was 488 nm supplied by an argon laser. The sperm population was gated using forward and side scatter. The volume of green (Annexin-V and Rhodamine-123) and red fluorescence were detected respectively with a FL1 photodetector (530 nm) and FL3 photodetector (610 nm). Next, 10×103 events were examined for each assay.

#### 2.10.1. Apoptosis status

For detection of sperm apoptosis status [25], the sperm samples were washed in calcium buffer and in next step, adding 10 μL Annexin V FITC (AV) was performed. Following incubating for a minimum of 20 min, 10 μL of propidium iodide (PI) was added to sperm suspension, then incubated for 10 min before flow cytometry. Following flow cytometry, sperm subpopulations process were classified into four various groups including: (1) viable non-apoptotic (AV-/PI-); (2) early apoptotic (AV+/PI-); (3) late apoptotic (AV+/PI-); and (4) necrotic (AV-/PI-) cells (Fig. 1). The late apoptotic and necrotic sperm were classified as dead sperm.

**Fig. 1.**
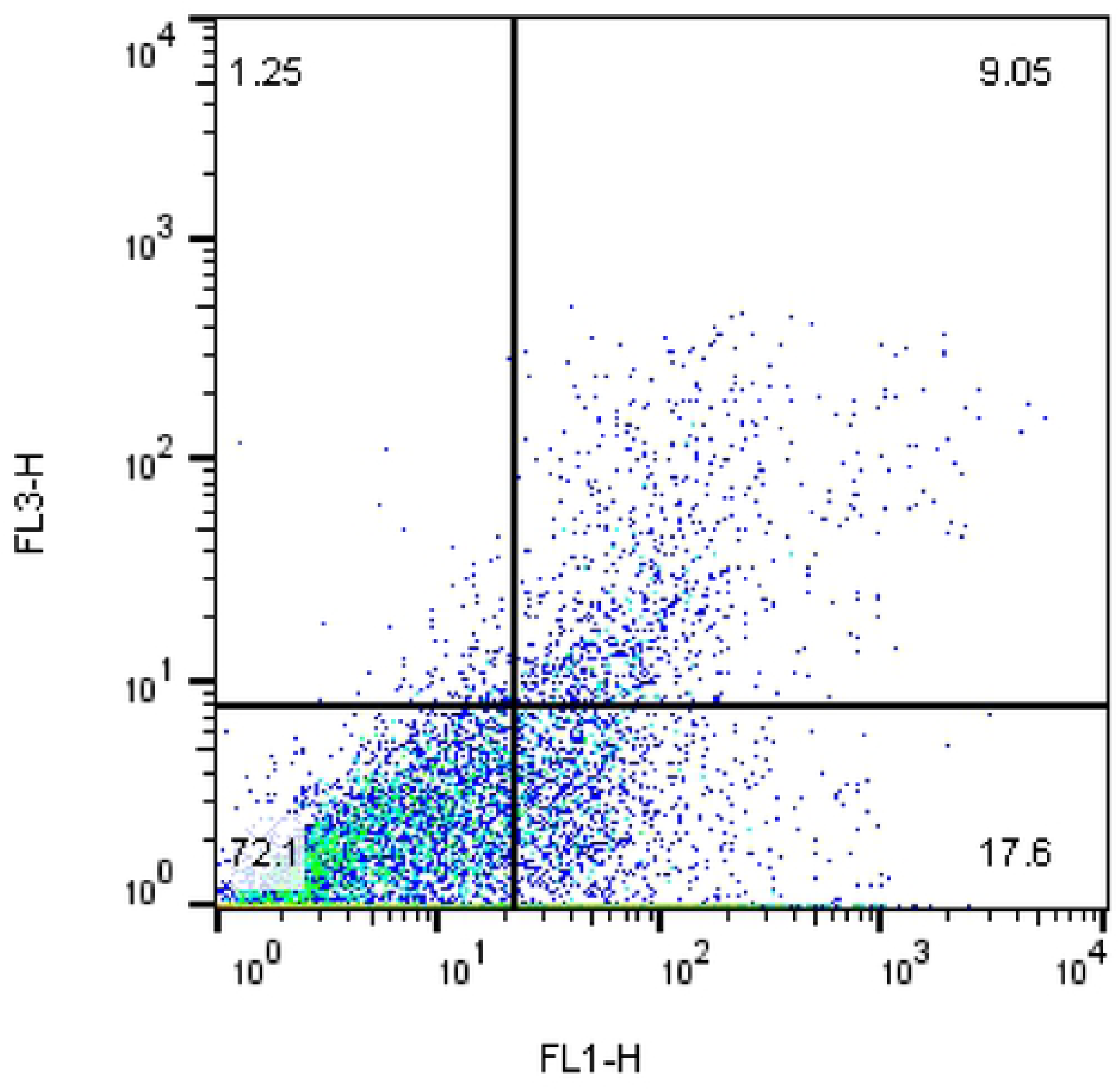
Annexin V and propidium iodide staining were used to determine the different cell populations.

#### 2.10.2. Mitochondrial activity

Mitochondrial activity was assessed by Rhodamine 123 (R123) and PI staining [26]. In brief, 5 microliters of R123 solution (0.01 mg/ml) and PI were added to 250 µl of diluted semen sample and, then incubated in dark place for 20 min. At last, the percentage of sperm mitochondrial activity (positive signal for Rh123 and negative signal for PI) was assessed by flow cytometer (Fig. 2).

**Fig. 2.**
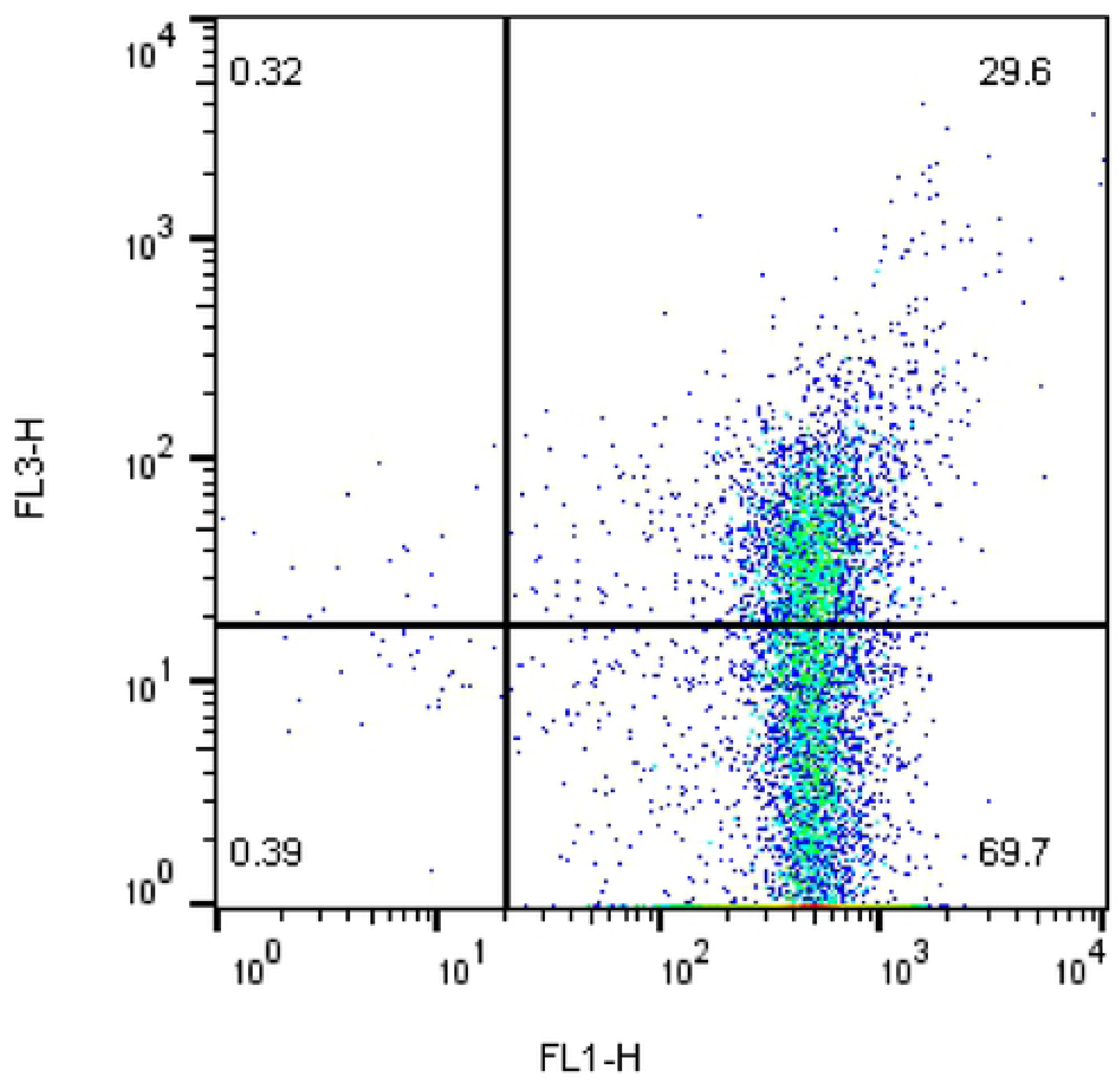
Flow cytometric detection of rooster sperm stained with Rhodamine123 and PI after freeze-thaw process.

### 2.11. RNA extraction and real-time polymerase chain reaction

Primers were designed using Primer3Plus online software on the basis of GenBank sequence of target genes and are presented in Table 2. The specificity of the primers was checked by a BLAST analysis of the National Center for Biotechnology information’s database. At the meantime, GAPDH was amplified as an endogenous control gene.

**Table 2.**
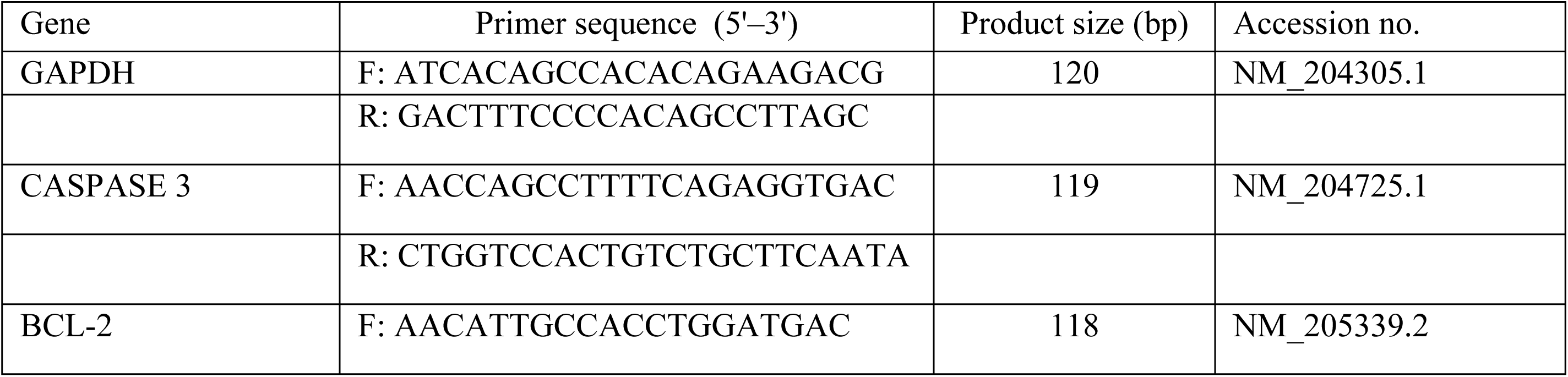
Primer sequences used for quantitative real-time polymerase chain reaction

Total RNA was extracted from sperm samples using Trizol reagent (Invitrogen, Carlsbad, CA, USA) following the method provided by the manufacturer and quantified using ND-1000 spectrophotometer (NanoDrop Technologies, Wilmington, DE, USA). RNA was transcribed into complementary DNA with the reverse transcription reagent kit (REVERTA-L RT reagents kit; code: K3-4-100-CE) and a thermal cycler according to manufacturer’s instructions. The RT reaction was conducted in 20 mL of reaction mixture at 37 °C for 15 minutes and then stored at ≤ –20 °C.

All polymerase chain reactions (PCRs) were carried out in ABI StepOnePlus Real-Time PCR Systems (Applied Biosystems, USA) using the RealQ Plus 2x Master Mix Green Kit (Ampliqon, code: A325402) following manufacturer’s instructions. As a whole, the reaction was performed at 95 °C for 15 min, followed by 40 cycles of denaturing, and annealing and elongating (95 °C for 15 seconds, 61 °C for 20 seconds and 72 °C for 30 seconds, respectively). The dissociation curves of PCR products were achieved by a following cycle of 95 °C for 15 seconds, 60 °C for 1 min and 95 °C for 15 seconds, and reaction specificity was defined when there was only one specific peak in the dissociation curve. The R2 values for all standard curves generated ranged 0.999, and PCR efficiencies was ≥95%. The quantitative PCR data were analysed using the 2-ΔΔCt method (Livak and Schmittgen 2001).

### 2.12. Artificial insemination

Reproductive performance of post-thawed sperm was assessed by artificial insemination [3]. A total of 30 Ross broiler breeder hens were caged (10 hens in each group) and fed a standard diet. The straws from each treatment were thawed and inseminated with a dose of 100 × 106 sperm. Eggs were collected for five days after the last artificial insemination. The eggs were incubated in a commercial incubator. On day 7 of incubation, fertility (by candling the eggs) and hatchability (the percentage of hatched eggs per fertile eggs) were evaluated for each treatment.

### 2.13. Statistical analysis

Data obtained from post-thawing quality were analyzed by PROC GLM SAS 9.1 (version 9.1, 2002, USA). Effects of supplemented antioxidant on fertility and hatchability were analyzed using GENMOD procedure. The results are expressed as the mean±SEM. The Turkey’s test was performed to compare treatments. Significance level was adjusted to p < 0.05.

## 3. Results

Motility and velocity variables of frozen-thawed of rooster sperm supplemented with different levels of crocin and naringenin are depicted in Table 3. C1 and N100 resulted in higher (P < 0.05) total motility and progressive motility compared to the control group. The analysis did not reveal any significant differences among different concentrations of crocin and naringenin on the VCL, VAP, VSL, ALH, LIN, BCF and STR parameters.

**Table 3.** Effect of different levels of crocin and naringenin on motility parameters of rooster thawed semen, analyzed by CASA (*n* = 5).

| Antioxidant | TM (%) | PM (%) | VSL (μm/s) | VAP (μm/s) | VCL (μm/s) | LIN (%) | STR (%) | ALH (μm) | BCF (Hz) |
| --- | --- | --- | --- | --- | --- | --- | --- | --- | --- |
| Control | 61.06<sup>c</sup> | 22.03<sup>b</sup> | 16.37 | 29.94 | 52.47 | 32.88 | 54.65 | 5.25 | 15.33 |
| C0.5 | 65.19<sup>bc</sup> | 26.66<sup>ab</sup> | 17.42 | 31.74 | 56.22 | 31.32 | 54.80 | 4.94 | 15.72 |
| C1 | 74.43<sup>a</sup> | 30.88<sup>a</sup> | 18.78 | 33.13 | 57.72 | 33.03 | 57.88 | 4.58 | 16.10 |
| C1.5 | 61.34<sup>c</sup> | 25.10<sup>ab</sup> | 16.76 | 30.42 | 54.08 | 31.28 | 55.86 | 4.90 | 15.17 |
| N50 | 64.13<sup>bc</sup> | 24.58<sup>ab</sup> | 17.30 | 31.24 | 53.23 | 32.14 | 56.51 | 5.28 | 15.59 |
| N100 | 71.21<sup>ab</sup> | 28.46<sup>a</sup> | 18.68 | 32.01 | 57.13 | 33.02 | 59.25 | 4.78 | 16.06 |
| N150 | 60.94<sup>c</sup> | 21.61<sup>b</sup> | 16.54 | 30.33 | 51.08 | 31.26 | 55.59 | 5.04 | 15.49 |
| SEM | 1.78 | 1.79 | 1.63 | 1.96 | 2.55 | 3.68 | 6.07 | 0.26 | 1.77 |

MOT: Total motility (MOT, %); PROG: Progressive motility; VSL: straight-line velocity; VAP: Average path velocity; VCL: curvilinear velocity; LIN: Linearity; STR: Straightness; ALH: Mean amplitude of the lateral head displacement; BCF: Mean of the beat cross frequency. Different superscripts within the same column indicate significant differences among groups ( $p < 0.05$ ).

The findings of the current research revealed that plasma membrane integrity in C1 and N100 were significantly higher compared to control group (Fig. 3). Fig. 4 summarizes the data on mitochondrial activity. The findings of this test revealed that the percentage of mitochondria activity was higher in the C1 and N100 groups. The results show that different levels of crocin and naringenin does not seem to impact the abnormal forms after freeze-thawing (Fig. 5). Superior results were observed for viable sperm in C1 and N100 compared with control group (Fig. 6).

**Fig. 3.**
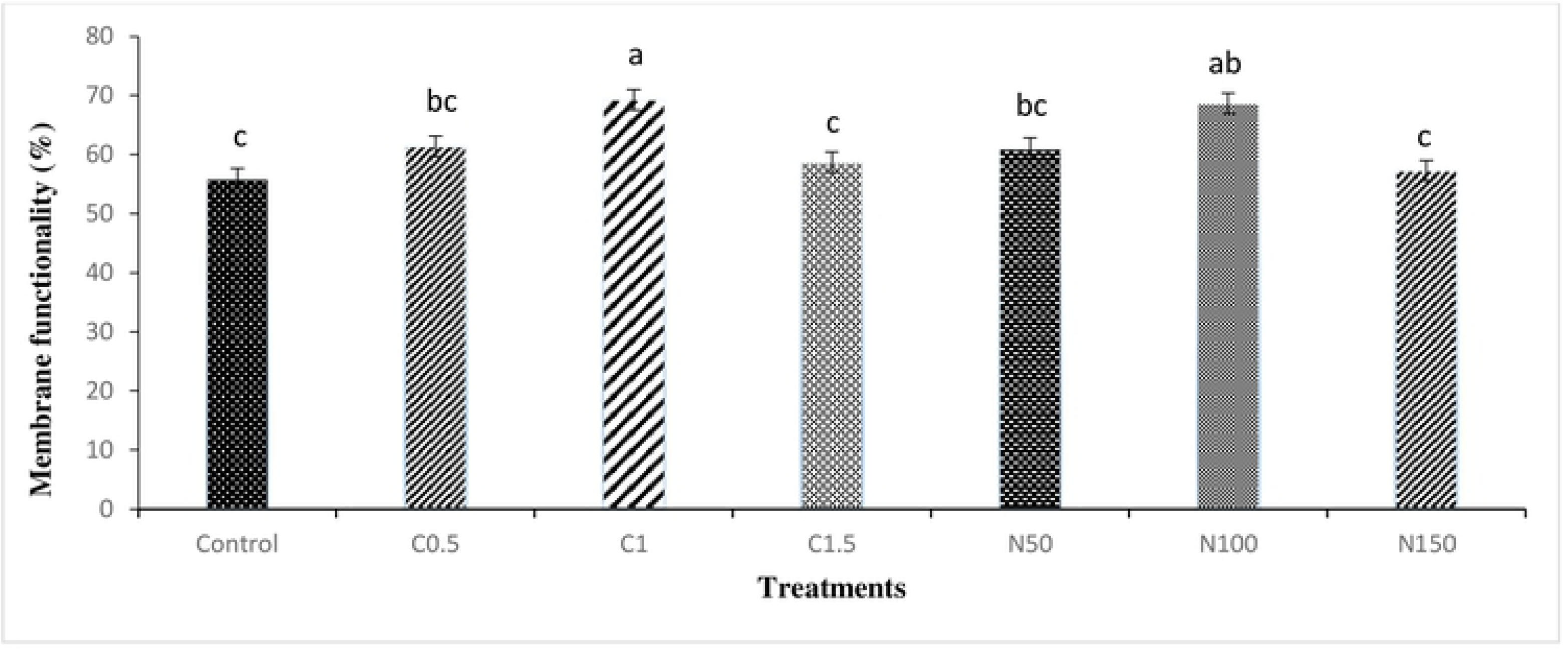
Effect of crocin and naringenin supplementation in cryopreservation medium on post-thawed membrane functionality of rooster sperm. Beltsville extender without antioxidant (control), C0.5 (Beltsville extender with 0.5 mM crocin), C1 (Beltsville extender with 1 mM crocin), C1.5 (Beltsville extender with 1.5 mM crocin), N50 (Beltsvil1e extender with 50 µM naringe nin), N100 (Beltsville extender with 100µM naringenin), N150 (Beltsville extender with 150 µM naringenin).

**Fig. 4.**
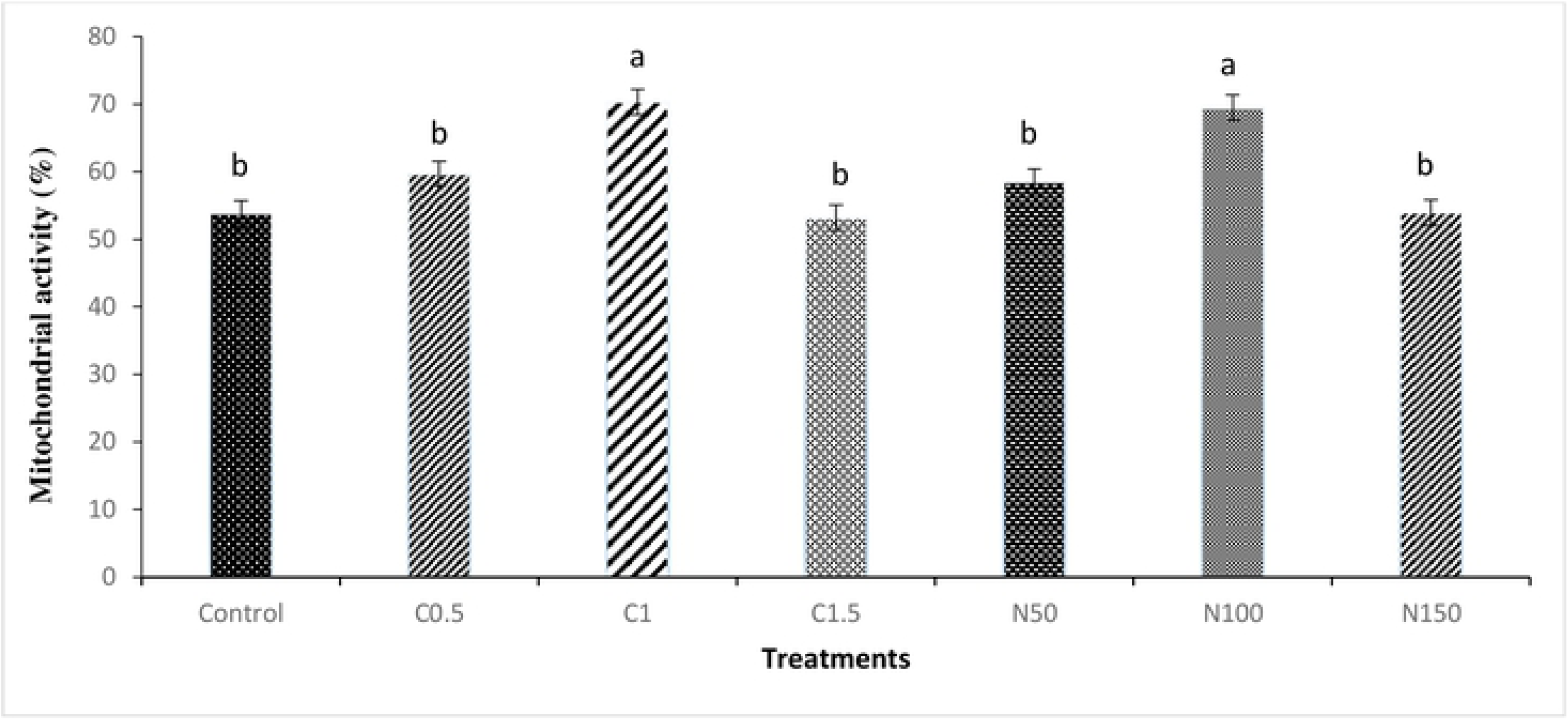
Effect of crocin and naringenin s upplementation in cryopreservation medium on post-thawed mitochondrial activity of rooster sperm. Beltsville extender without antioxidant (control), C0.5 (Beltsville extender with 0.5 mM crocin), C1 (Beltsville extender with 1 mM crocin), C1.5 (Beltsville extender with 1.5 mMcrocin), N50 (Beltsville extender with 50 **µM** naringenin), N100 (Beltsville extender with 100 **µM** naringenin), N150 (Beltsville extender with 150**µM** naringenin).

**Fig. 5.**
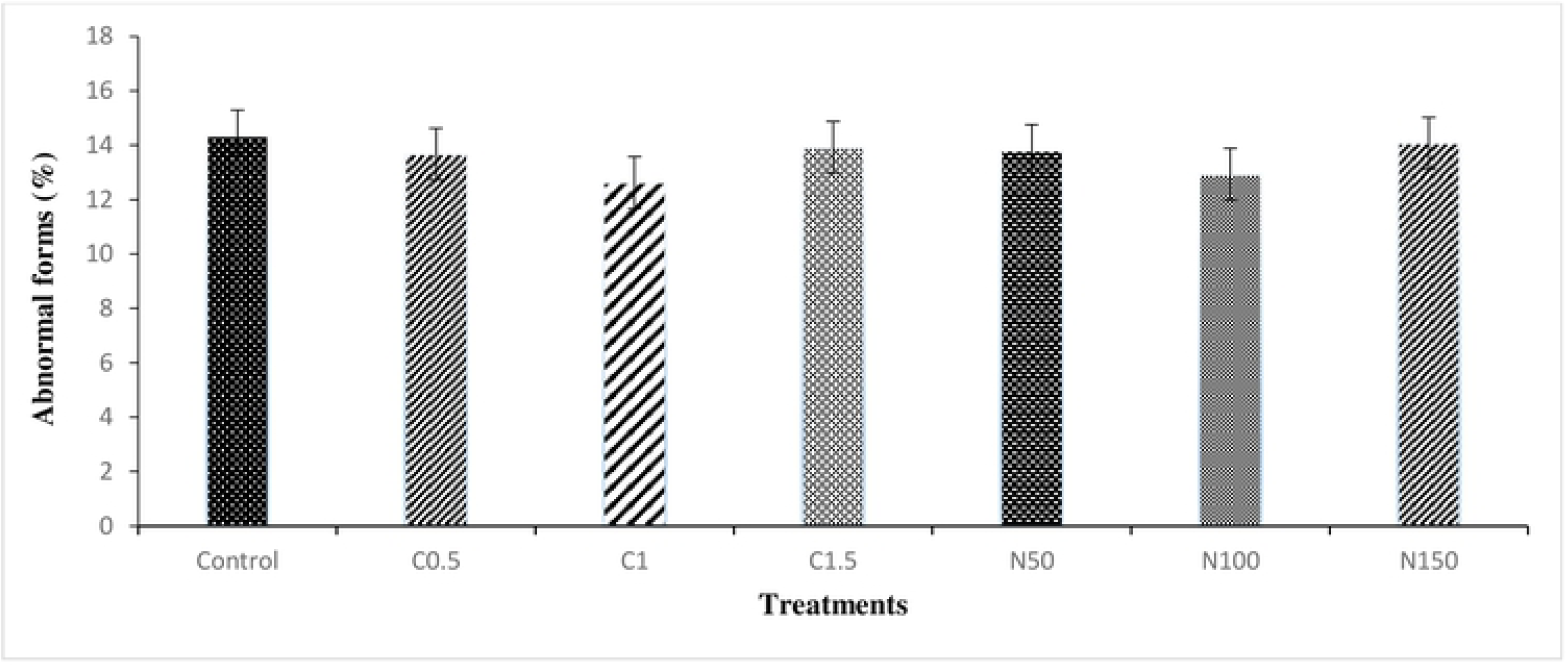
Effect of crocin and naringenin s upple mentation in cryopreservation medium on post-thawed abnormal forms of rooster sperm. Beltsville extender without antioxidant (control), C0.5 (Beltsville extender with 0.5 mM crocin), C1 (Beltsville extender with I mM crocin), C1.5 (Beltsville extender with 1.5 mMcrocin), N50 (Beltsville extender with 50 µM naringc nin), N100 (Beltsville extender with 100 µM naringcnin), N150 (Beltsville extender with 150 µM naringenin).

**Fig. 6.**
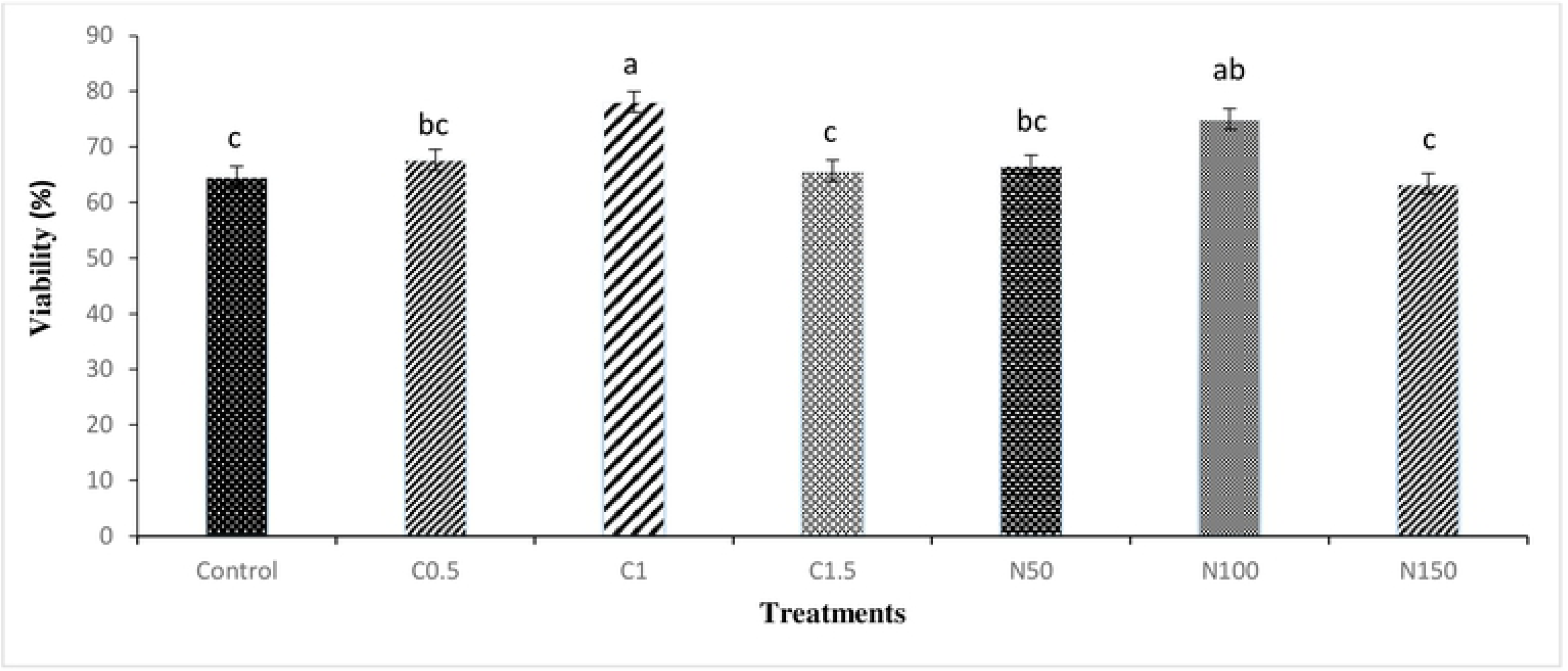
Effect of crocin and naringenin supplementation in cryopreservation medium on post-thawed viability of rooster sperm. Beltsville extender without antioxidant (control), C0.5 (Beltsville extender with 0.5 mM crocin), C1 (Beltsville extender with 1 mM crocin), C1.5 (Beltsville extender with 1.5 mM crocin), N50 (Beltsville extender with 50 µM naringenin), N100 (Beltsville extender with 100 µM naringenin), N150 (Beltsville extender with 150 µM naringenin).

Table 4 details the data on apoptosis status analysis. The most remarkable result is that the percentage of live sperm was emerged to be higher in 1 mM crocin and 100 µM naringenin in comparison with the control. Apoptotic spermatozoa were significantly reduced in the C1 and N100 levels when compared to control group.

**Table 4.** Effect of different levels of crocin and naringenin on viable, apoptotic and dead sperm in rooster thawed semen, as assessed by flow cytometry (*n* = 5).

| Antioxidant | Live (%) | Early apoptosis (%) | Dead (%) |
| --- | --- | --- | --- |
| Control | 56.95<sup>b</sup> | 25.28<sup>a</sup> | 17.76 |
| C0.5 | 60.88<sup>b</sup> | 21.88<sup>ab</sup> | 17.23 |
| C1 | 71.46<sup>a</sup> | 15.30<sup>b</sup> | 13.23 |
| C1.5 | 57.31<sup>b</sup> | 24.47<sup>a</sup> | 18.20 |
| N50 | 59.73<sup>b</sup> | 22.81<sup>ab</sup> | 17.45 |
| N100 | 70.86<sup>a</sup> | 15.40<sup>b</sup> | 13.72 |
| N150 | 57.24<sup>b</sup> | 24.78<sup>a</sup> | 17.97 |
| SEM | 1.39 | 1.81 | 1.87 |

Different superscripts within the same column indicate significant differences among groups ( $p < 0.05$ ). Beltsville extender without antioxidant (control), C0.5 (Beltsville extender with 0.5 mM crocin), C1 (Beltsville extender with 1 mM crocin), C1.5 (Beltsville extender with 1.5 mM crocin), N50 (Beltsville extender with 50 μM naringenin), N100 (Beltsville extender with 100 μM naringenin), N150 (Beltsville extender with 150 μM naringenin)

Table 5 reports the data on effects of various levels of crocin and naringenin on the oxidative parameters status of rooster sperm following freeze-thawing. We can note from the table that the highest values for TAC activity were achieved in the C1 and N100 groups compared with control group. Also, malondialdehyde was significantly (P < 0.05) lower in C1 and N100 than the control. The analysis did not reveal any significant differences for SOD and GPx parameters.

**Table 5.** Effect of different levels of crocin and naringenin on malondialdehyde concentration (MDA), glutathione peroxidase (GPx) and superoxide dismutase (SOD) activities and total antioxidant capacity (TAC) of rooster thawed semen (n = 5).

| Antioxidant | MDA<br>(nmol/mL) | GPx<br>(U/mg protein) | SOD<br>(U/mg) | TAC<br>(mmol/l) |
| --- | --- | --- | --- | --- |
| Control | 4.26<sup>a</sup> | 54.00 | 107.70 | 1.13<sup>bc</sup> |
| C0.5 | 2.91<sup>c</sup> | 60.70 | 118.25 | 1.70<sup>ab</sup> |
| C1 | 1.83<sup>d</sup> | 63.20 | 124.65 | 1.85<sup>a</sup> |
| C1.5 | 3.81<sup>ab</sup> | 55.20 | 109.38 | 1.26<sup>c</sup> |
| N50 | 3.12<sup>bc</sup> | 57.85 | 117.87 | 1.66<sup>abc</sup> |
| N100 | 1.90<sup>d</sup> | 62.71 | 123.92 | 1.88<sup>a</sup> |
| N150 | 4.01<sup>a</sup> | 53.61 | 108.37 | 1.18<sup>c</sup> |
| SEM | 0.19 | 2.14 | 6.33 | 0.13 |

Different superscripts within the same column indicate significant differences among groups ( $P < 0.05$ ). Beltsville extender without antioxidant (control), C0.5 (Beltsville extender with 0.5 mM crocin), C1 (Beltsville extender with 1 mM crocin), C1.5 (Beltsville extender with 1.5 mM crocin), N50 (Beltsville extender with 50 μM naringenin), N100 (Beltsville extender with 100 μM naringenin), N150 (Beltsville extender with 150 μM naringenin)

The results of mRNA expressions of BCL-2 and CASPASE 3 are showed in Fig.7 and Fig. 8. The mRNA expressions of BCL-2 in the C1 and N100 group were significantly higher than that in other treatments. The expression of CASPASES 3 was significantly reduced in C1 and N100 group (P < 0.05) when compared to control group.

**Fig. 7.**
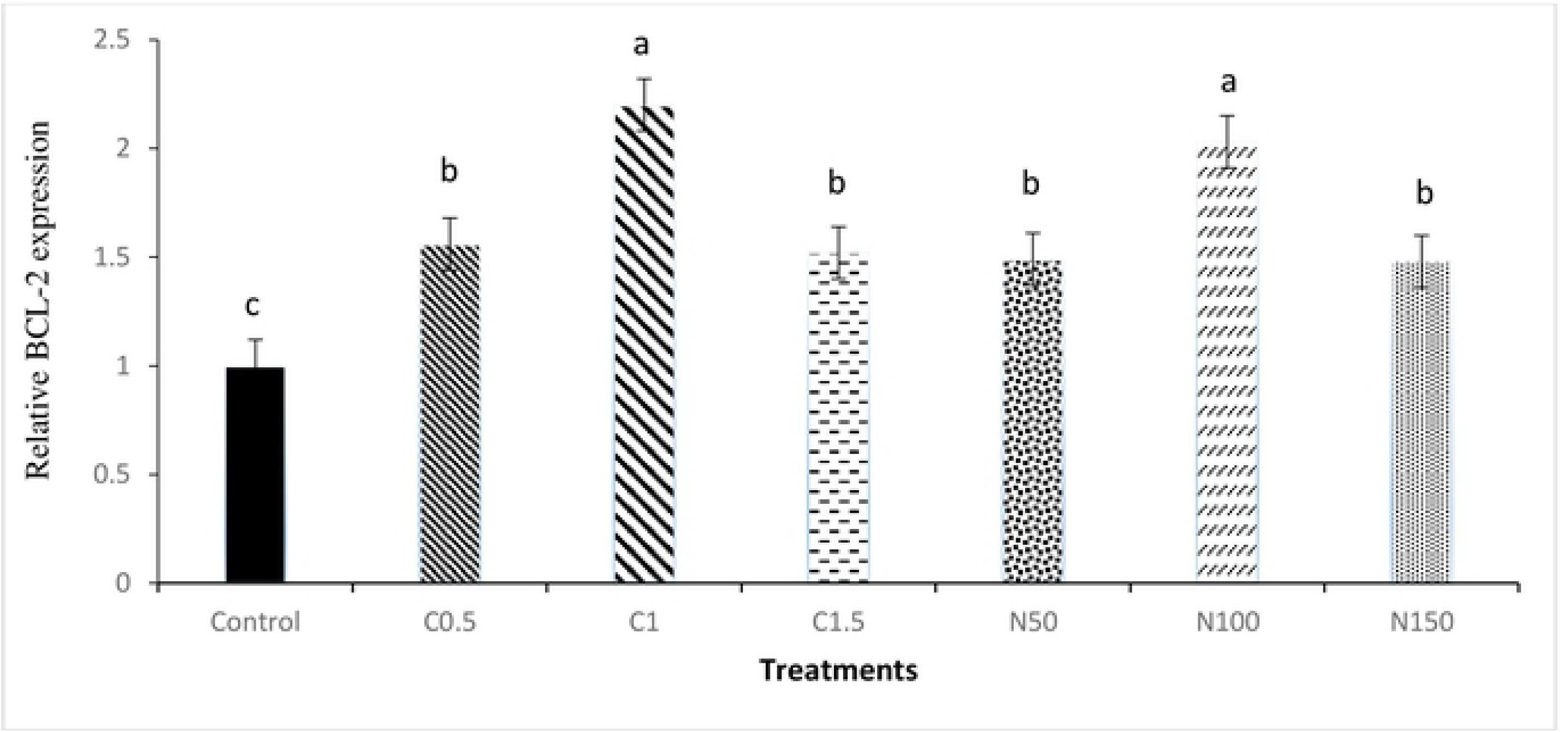
Relative mRNA expression of BCL-2 gene in therooster sperm cryopreservation. Beltsville extender without antioxidant (control), C0.5 (Beltsville extender with 0.5 mM crocin), C1 (Beltsville extender with **1** mM crocin), C1.5 (Beltsville extender with 1.5 **mM** crocin), N50 (Beltsville extender with 50 **µM** naringenin), N1OO(Beltsville extender with 100 **µM** naringenin), N150 (Beltsville extender with 150 **µM** naringenin).

**Fig. 8.**
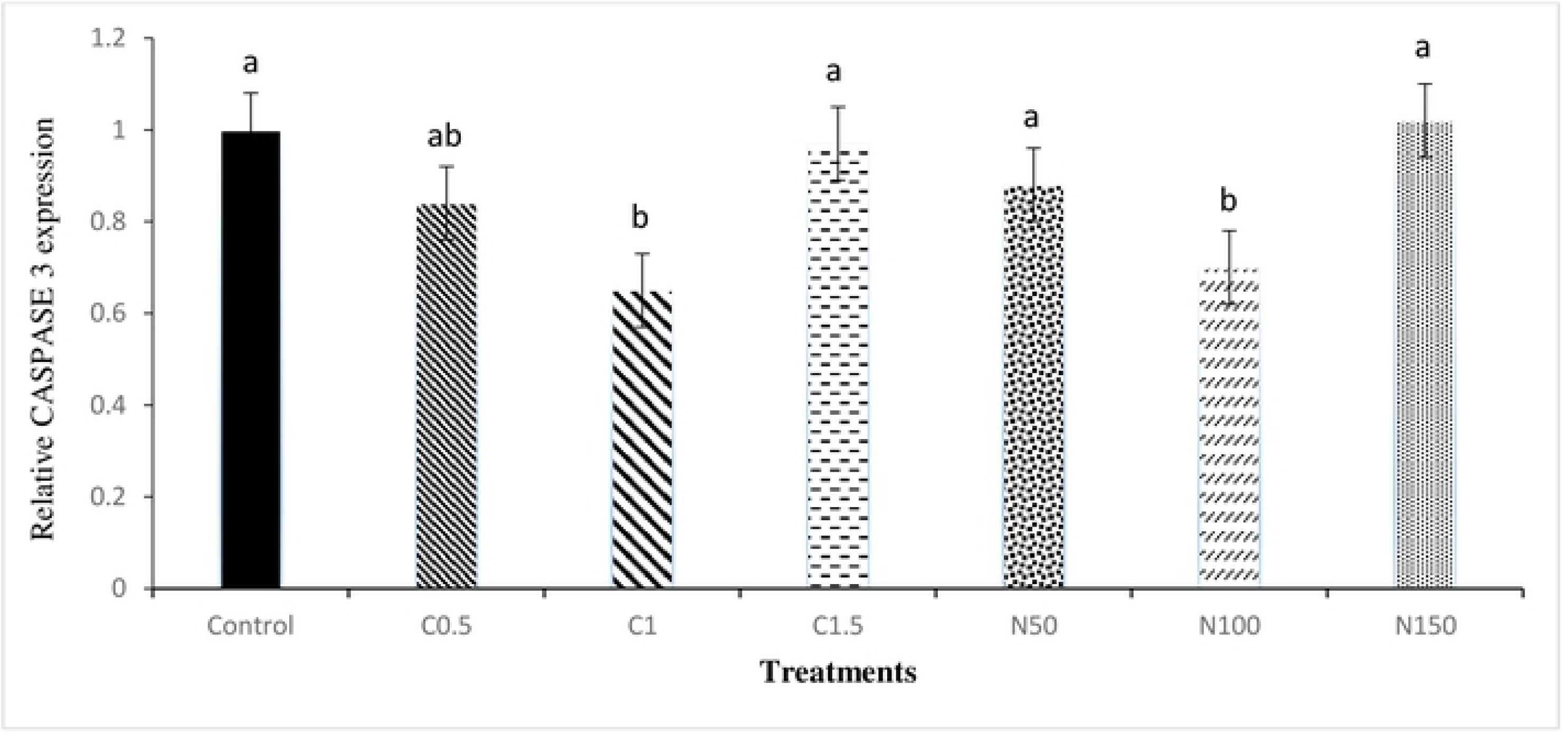
Relative mRNA expression of CASPASE 3 genein the rooster sperm cryopreservation. Beltsville extender without antioxidant (control). C0.5 (Beltsville extender with 0.5 mM crocin), C1 (Beltsville extender with I mM crocin), C1.5 (Beltsville extender with 1.5 mM crocin), N50 (Beltsville extender with 50 µM naringenin), N1OO(Be ltsville extender with 100µM naringenin), N150 (Beltsville extender with 150 µM naringenin).

The findings of the fertility trial (Table 6) revealed a significantly higher (P < 0.05) percentage of fertility and hatching rate in C1 and N100 compared to the control group.

**Table 6.** Effect of crocin and naringenin on fertility and hatchability rates of rooster semen after freeze-thawing

| Antioxidant | Fertility (%) | Hatchability of fertile egg (%) |
| --- | --- | --- |
| Control | 36<sup>b</sup> | 47.37<sup>a</sup> |
| Naringenin 100 µM | 54<sup>a</sup> | 74.07<sup>a</sup> |
| Crocin 1 mM | 52<sup>a</sup> | 73.08<sup>a</sup> |

Different superscripts letters within column are significantly different ( $P < 0.05$ ).

## 4. Discussion

Studies evaluating the efficacy of antioxidants to prevent damage during sperm cryopreservation usually cause contradictory results. Some experiments have reported a protective effect against cryo-related oxidative damages [27]. However, other studies could not show significant effects; some even led to impaired sperm function [10, 28, 29]. In this context, some points must be taken into consideration when performing an antioxidant treatment. An important point is that each ROS is deactivated by a specific antioxidant system [30, 31]; therefore, if antioxidant therapy is chosen at random, this treatment will not be effective if it is not directed to ROS, which is the main cause of oxidative damage [32]. The protective effect of natural antioxidants on oxidative stress in avian species has been considered in various studies. However, such antioxidants are generally present in little amounts in semen for neutralizing the oxidative stress which occurs during in vitro sperm conservation [33]. It is demonstrated that defeating oxidative stress can be supplied by adding a variety of antioxidants to the bird sperm [21, 23, 34]. Some characteristics of crocin and naringenin make them highly effective supplements to be utilized as additives for rooster sperm cryopreservation medium. The hypothesis in the present study was that crocin and naringenin, as a supplement in freezing extenders, could be effective in eliminating oxidative damage caused by freezing.

It was observed in our study that the addition of 1 mM of crocin and 100 µM naringenin during preparation of the sperm had a beneficial effect on the total and progressive motility of sperm in comparison with the control group, while no effect was observed on the other motility parameters. The favorable effect of saffron and its bioactive component, crocin, on some parameters such as motility and viability has been demonstrated in deer, mice and humans [35–37]. It is demonstrated that in stressful conditions, naringenin has the ability to chelate irons and decrease ROS production. Interestingly, it is related to the fact that naringenin has 5-hydroxy and 4-carbonyl groups in the C-ring which plays a role in ROS scavenging, Cu, and Fe ions interaction [38, 39]. Therefore, adding naringenin to the cryopreservation medium can decrease the stress caused by freezing, consequently can increase motility which was observed in our study.

It is shown that crocin can reduce the levels of superoxide anion and hydrogen peroxide. The supplementation of crocin in the cryopreservation medium showed to be advantageous for the sperm in terms of viability at the C1 group. Carotenoids show stabilizing effect on sperm conservation by interaction with the superoxide anion [40]. Furthermore, crocin enhances the activity of particular intracellular detoxifying enzymes or effects the fluidity of the membrane, which influences its permeability to oxygen and further molecules [41].

Our previous studies has adopted an approach in the study of the associations between sperm variables and MDA levels [42]. The correlation between MDA content of the sperm and the fertilization capacity is worth mentioning [43]. Malondialdehyde levels in semen are inversely proportional to the function of sperm [32, 44]. These data were again confirmed in the present investigation, in which the MDA level was evaluated because it is known as a gold marker for oxidative stress, a phenomenon extremely associated to the antioxidant system. In line with our study, Sapanidou, Taitzoglou (17), showed that MDA production decreased while supplementing 1 mM crocin in sperm.

According to our results, naringenin 100 µM reduced the MDA level. A satisfactory explanation for this may be related to its structure-activity. Naringenin can give hydrogen to ROS that allows the acquisition of a stable composition, allowing the elimination of these free radicals. Another interesting reason is the existence of phenolic rings in naringenin which act as electron barriers to remove superoxide anions characteristic known as free radicals [45].

In the light of the data, it is clearly essential to comprehend what cellular factors normally serve as causes for free radical production by mitochondria of sperm. It is demonstrated that the generation of mitochondrial ROS raises when the membrane potential collapses pharmacologically [46]. Carotenoids have a recognized protective effect in the mitochondria and the crocin itself has been reported a mitochondrial protector [47]. Therefore, it was predictable that C1 and N100 increased mitochondrial activity after thawing. The axosomas and dense fibers associated with the central part of the sperm cells are covered by mitochondria, the organs which produce energy from ATP that are involved in sperm motility [48]. It is obvious that cryopreservation results in a reduction in sperm motility, morphological functional integrity and mitochondrial membrane potential by inducing axonemal damage [49–51]. Sperm motility is the main key for the substantial penetration of cumulus cells and the zona pellucida of the ovum [48]. A conspicuous correlation is confirmed between sperm motility and mitochondrial activity [3]. Therefore, in the present study, supplementation of sperm extender with crocin 1mM and 100 µM naringenin before cryopreservation increased membrane integrity and mitochondrial activity leading to improving sperm motility.

Mitochondrial dysfunction is shown to be a critical modulator of ROS production and consequently onset of apoptosis. An interesting result was found for crocin 1mM and 100 µM naringenin in reducing early apoptosis. This is in complete agreement with Sapanidou et al. who reported that PS externalization decreased in the group containing 1mM crocin [17]. Our results do not support the observations by Mata-Campuzano, Alvarez-Rodriguez (52), who noted that crocin did not affect apoptotic ratio in ram sperm following cryopreservation. It is indicated that various apoptogenic proteins containing Cyt-c, AIF and Endo-G are released through pores generated by the mitochondrial membrane potential and consequently inhibiting the release of different types of apoptogenic factors from mitochondria. Thereby, the expressions of caspase-3 and bcl-2 which were regulated in sperm cells owing to the release of apoptogenic factors from mitochondrial pores was inhibited in naringenin 100 and crocin 1. As explained above, naringenin is effective in conserving the mitochondrial membrane by preventing the excessive production of ROS, consequently, inhibiting the release of several apoptogenic factors from the mitochondria [53]. Also it is shown that naringenin restricts translocation of AIF and Endo-G to the nucleus by restoring mitochondrial membrane potential that prevents DNA damage and, finally inhibits cell damage [54]. It is an appreciable reason for preventing apoptosis by naringenin after freeze thawing. Naringenin can initiate the mitochondrial-mediated apoptosis pathway as revealed by an enhanced ratio of (pro-apoptotic) Bax/(anti-apoptotic) Bcl2 genes, therefore results in release of cytochrome C and consequent activation of Caspase-3 [55]. Caspase-3 is known as the critical effector caspase responsible for the execution of apoptotic cell death by cleaving numerous cellular substrates [56].

The results of this study show that the enhancement in fertility result using thawed sperm stored in C1 and N100 was consistent with the other sperm functional parameters. The freezing and thawing process dramatically reduces the fertilization capacity of the rooster sperm. Likewise, a relatively large number of live sperm is required inside the sperm storage tubes (SST) to determine fertilization after inseminations [57]. The semen parameters related to fertility such as sperm motility, vitality and progressive motility can influence the penetration of cervical mucus. So, the strategies that enhance the sperm viability and motility will ensure sperm journey in the hen reproductive tract to attain SST and then the fertilization position. Also, enhancement of cellular variables by improving sperm antioxidant system and mitochondria activity will increase sperm function during passage in the reproductive tract [25]. Therefore, it appears that higher sperm quality in a group of C1 and N100 showed greater hatching among treatment groups by preserving more alive sperm in SST and influencing fertility in the current trial.

The present study showed that 1 mM crocin and 100 µM naringenin could beneficially affect a variety of semen quality in Ross 308 breeder roosters. Particularly, 1 mM crocin and 100 µM naringenin could protect the sperm against excessive ROS generation by reducing the pro-apoptotic (CASPASE 3) and increasing anti-apoptotic (Bcl-2) apoptosis genes. Also, enrichment of semen extender with 1 mM crocin and 100 µM naringenin improved fertilizing capacity of rooster sperm.

